# FlowerPatch: New Method to Measure Nectar Volume in Artificial Flowers

**DOI:** 10.1101/2025.02.16.638513

**Authors:** E. Lara, J. Agosto, G. Tugrul, R. Megret, E. Flórez

**Affiliations:** University of Puerto Rico Mayaguez; University of Puerto Rico Rio Piedras; University of Puerto Rico

## Abstract

This article proposes a new Flower Patch Nectar Sensor to address the problem of detecting and measuring nectar in artificial flowers used in experiments on pollinator behavior. Traditional methods have focused mainly on recording the visits of pollinators to the flowers, without addressing the dynamic variations in nectar in terms of volume and concentration. The proposed approach provides more detailed information about the nectar consumption by bees and allows determination of the optimal time to refill the flowers. This study introduces an innovative method that uses electrodes and an oscillator circuit to measure the volume of nectar present in the flower. The system correlates the concentration of nectar with a frequency signal that can be processed by a microcontroller. It was evaluated using initial volumes ranging from 1 µL to 4 µL, demonstrating its ability to accurately detect variations in nectar, even up to the point where the frequency approaches zero. The results confirm that this method allows us to identify how the reward offered to pollinators (represented by nectar) varies over time, in terms of concentration, under both controlled and natural conditions. Additionally, graphs are presented that show the relationship between an initial volume of 4 µL and variations in the frequency signal over a period of 25 minutes, highlighting the influence of these factors on nectar dynamics. This work not only introduces an innovative approach for the dynamic monitoring of nectar in artificial flowers, but also lays the groundwork for future studies on the physical and chemical modeling of nectar in response to environmental conditions.

## Introduction

The study of the behavior of honey bees in their interaction with artificial flowers is essential to understand foraging patterns and the dynamics of nectar collection. However, most previous research has focused solely on measuring the number of visits to flowers, without addressing the variables that affect the amount of nectar available in them. For example, Kuusela and Lamsa (2016) and Debeuckelaere et al. (2022), research that focuses on the automation of nectar delivery and data collection, but does not include nectar detection, limiting itself to delivery and monitoring of visits. The systems of Essenberg (2015) and Sokolowski and Abramson (2010) focus on artificial flowers that deliver nectar in a discrete or controlled manner, but do not control or detect the amount of nectar delivered. Others, such as Keasar (2000) and Moffatt (2001), manipulate variables such as reward rate, spatial arrangement of flowers or metabolic conditions, but again, do not use sensors to measure or detect nectar.

In this context, the present work stands out by proposing an innovative methodology that not only measures visits, but also the volume of nectar remaining in artificial flowers. This allows for more precise data on bee behavior and nectar dynamics, which represents a significant advance in the understanding of nectar collection processes.

Furthermore, this study introduces a model that addresses how nectar concentration varies under different environmental conditions, specifically as a function of temperature and humidity. The model presented in this work establishes a precise relationship between the volume of nectar present and the current measured by the sensors, allowing the evaluation over time of the dynamic conditions that affect the nectar. The visual analysis that accompanies this model provides a clear and understandable representation of changes in nectar concentration, which is a valuable tool for future studies in the field.

The central innovation of this project lies in the use of a sensor that allows quantitative measurement of the presence and amount of nectar in artificial flowers. This approach, unlike previous studies that are mainly based on detecting the bee visit, contributes a new way of collecting data on nectar collection, contributing to the precision and reliability of the experimental results.

## System Architecture and Design

### General Architecture

The nectar sensing system is composed of three key mechanisms: a Flower Patch Nectar Sensor (FPNS) that measures the amount of nectar in the nectary with a 555 oscillator circuit that converts the variation of the intrinsic resistance of the nectar into a frequency signal, and a microcontroller (such as the Arduino UNO) that interprets the signal and controls the detection process. This system focuses solely on the precise measurement of nectar and its variation over time (See Figure 1).

**Figure 1.**
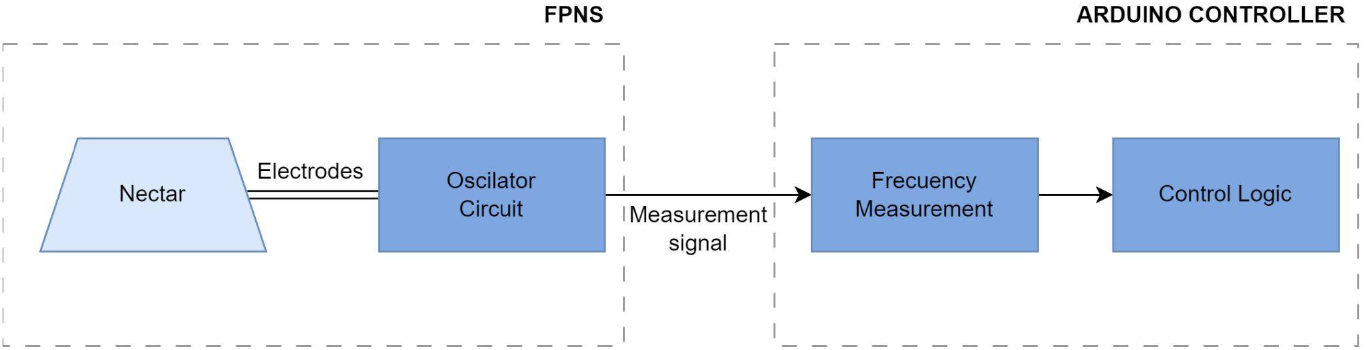
Block diagram of the nectar detection system correlating the volume of nectar with the output frequency of the oscillator circuit.

### Sensor design

The system is made up of the following essential components:

#### Flower Patch Nectar Sensor (FPNS)

This sensor is integrated into the artificial flower with electrodes at the bottom of the nectary coming into contact with the nectar. It also detects the intrinsic resistance of the nectar, and allows the conversion of this resistance into an output frequency using a 555 oscillator.

#### 555 Oscillator and Detection Circuit

The 555 oscillator is configured in astable mode and acts as a pulse generator. The frequency generated is proportional to the resistance of the nectar present in the sensor. This circuit is designed with external resistors and capacitors to adjust the duty cycle and output frequency (See Figure 2).

**Figure 2.**
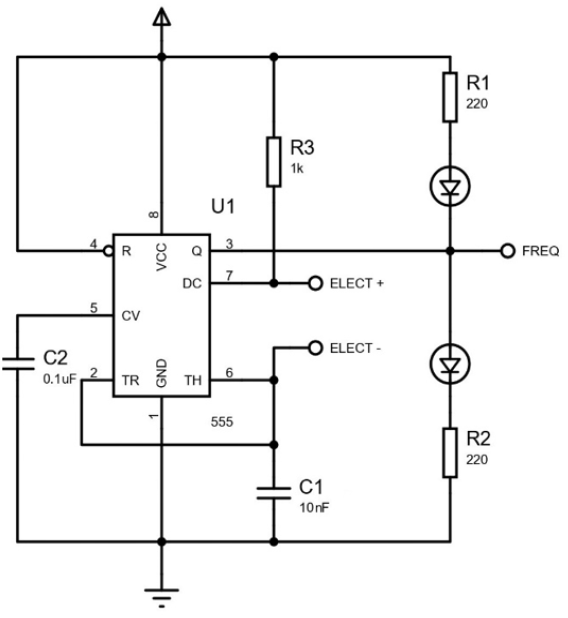
Oscillator Circuit Diagram for Nectar Detection. The circuit mainly uses a 1kΩ resistor, two capacitors, one of 0.1uF and one of 10nF to ensure an output frequency of 200 Hz for a 4uL nectar drop with an initial concentration of 2 M (molar) using 69.45 g of sugar in 100 mL of water.

#### Microcontroller and Algorithm

Was used a Arduino Uno to measure the frequency generated by the oscillator 555. AndThis microcontroller offers a simple and efficient way to capture these signals and send them for analysis. Frequency measurement is carried out through the pin 2, which is a interrupt pin, this allows detecting state changes in the signal without depending on constant queries within the code, and thanks to the fact that the data can be transmitted by serial communication, it is possible to store the data for later processing.

This is achieved with an arduino configuration for pin 2 as an external interrupt to detect state changes in the signal generated by the oscillator. 555, thus measuring its frequency. Every time a rising edge occurs, a counter is incremented and, after a time interval determined with the function millis(), the frequency is calculated by dividing the number of pulses by the elapsed time.

### Flower design

The design of the artificial flower used is composed of a 3D printed block that forms the accessible part of the flower, with a nectary located in the center and a side tube through which the nectar is supplied. At the base of the flower are the electrodes passing through the nectary, maintaining direct contact with the nectar for measurement (See Figure 3).

**Figure 3.**
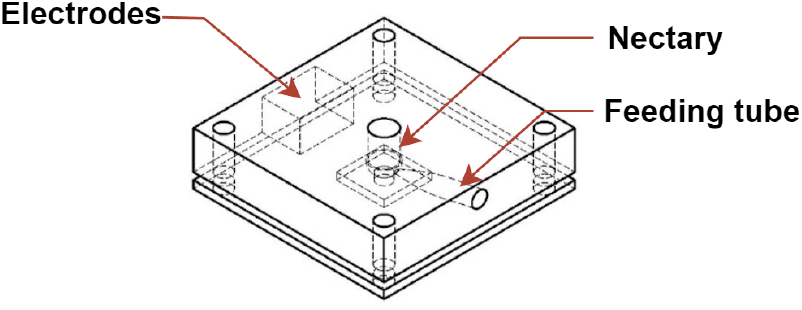
Artificial flower: Design with PCB and lead-free electrodes, lateral wiring maintaining a flat nectary.

## Experimental validation

Experimental validation of the Flower Patch Nectar Sensor (FPNS) system focused on evaluating its ability to detect changes in nectar concentration and volume in artificial flowers under controlled conditions (31°C y 61% RH). This sensor capable of directly measuring the signal coming from the nectar, Unlike those used to date, it proposes as an improvement to use a sensor based on a 555 oscillator circuit, capable of converting variations in the intrinsic resistance of the nectar into frequency signals that can be processed directly on the nectar solution.

In this section, two main experiments are presented. The first evaluates the sensor response over a 25-minute period, showing how evaporation affects frequency measurements. The second experiment analyzes the sensitivity of the FPNS to gradual increases in nectar, from 1 to 4 microliters, confirming its precision in detecting small variations in volume. These results not only validate the effectiveness of the system, but also establish a solid foundation for future research in controlled and natural environments.

The first graph (Figure 4) illustrates the behavior of the Flower Patch Nectar Sensor (FPNS) over a period of 25 minutes, recording changes in frequency in response to the initial application of 4 microliters of nectar. The data show a gradual decrease in frequency over time, which can be explained by the natural evaporation of nectar on the sensor surface. This phenomenon highlights the sensitivity of the FPNS to environmental conditions, such as temperature and humidity, which accelerate the evaporation process and affect the sensor measurements.

**Figure 4.**
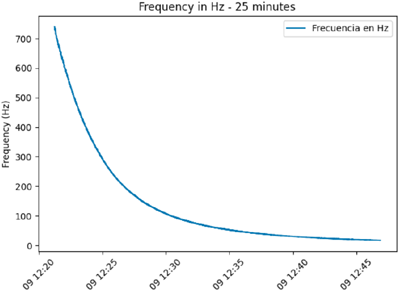
Nectar frequency over time

This decrease in frequency provides valuable information for behavioral experiments with bees, since it allows the volume and concentration of nectar to be estimated in real time. And by integrating computational methods, the system can dynamically adjust nectar availability, optimizing the reward system for pollinators. This ability is particularly relevant in natural environments where variables such as temperature fluctuations and direct sun exposure directly affect nectar solutions.

The second graph (Figure 5) shows the response of the sensor to the incremental addition of nectar, starting with 1 microliter and increasing by 1 microliter every 30 seconds until reaching 4 microliters. The frequency output exhibits a stepped pattern, reflecting the relationship between nectar volume and the frequency response of the sensor. Each step represents a discrete addition of nectar, with the sensor consistently recording higher frequencies for larger volumes.

**Figure 5.**
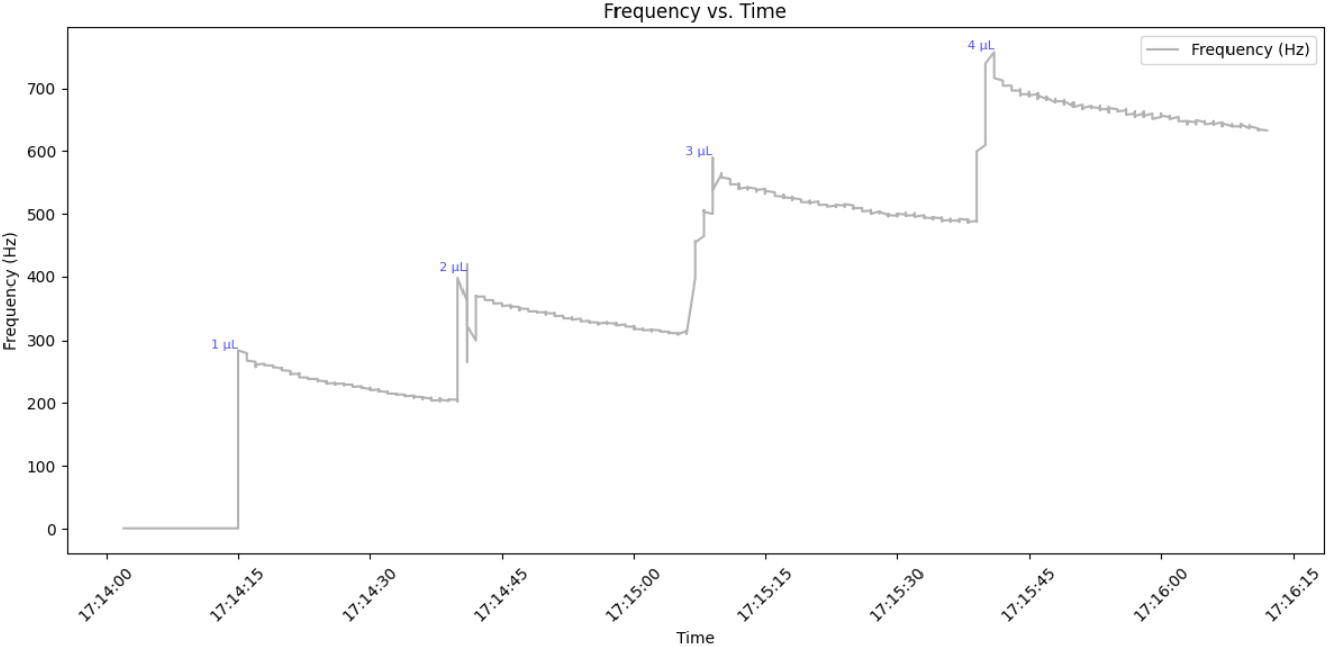
frequency changes with changes of 1 to 4 microliters of nectar

This stepwise behavior highlights the sensor’s ability to detect small variations in nectar volume accurately. Such sensitivity is crucial for experiments that seek to replicate natural conditions, where nectar availability can vary dynamically. By harnessing this capability, researchers can simulate diverse ecological scenarios, providing insight into the adaptability and decision-making processes of pollinators. Furthermore, the accuracy of the system supports the development of more accurate models of nectar dynamics, contributing to a deeper understanding of pollinator-plant interactions.

These results open new opportunities to carry out more complete and robust experiments, such as the integration with the work of Remi Megret, allowing to dynamically relate variables such as the amount of nectar delivered, and the approach to detecting the individual behavior of bees using convolutional neural networks ( Megret, 2024).

This provides a more comprehensive framework to analyze how controlled rewards, such as nectar, influence the behavior of marked bees under different experimental settings.

## Conclusions

The Flower Patch Nectar Sensor (FPNS) system demonstrated to be a new sensitive tool to dynamically measure the volume and concentration of nectar in artificial flowers, validating its ability to detect gradual variations and changes caused by evaporation under controlled conditions. The results highlight how environmental factors, such as temperature and humidity, impact nectar availability, underscoring the importance of integrating additional sensors to model and predict these dynamics. This approach not only allows for more precisely replicating natural scenarios, but also optimizes the rewards offered to pollinators by dynamically adjusting nectar availability. The capacity of the FPNS opens new possibilities to investigate the interaction between pollinators and flowers, as well as to design more efficient and controlled experiments. These findings constitute a solid basis for future research that seeks to understand how dynamic changes in nectar affect the behavior of pollinators and their relationship with the environment, contributing to the advancement of neuroscience.

